# BIOME-Preserve: A Novel Storage and Transport Medium for Preserving Anaerobic Microbiota Samples for Culture Recovery

**DOI:** 10.1101/2020.12.07.415638

**Authors:** Embriette R. Hyde, Hiram Lozano, Steven Cox

## Abstract

Culture-based study design is critical to advance research into the relationship between human health and the microbiome. Traditional sample collection protocols are focused on preserving nucleic acids and metabolites and are largely inappropriate for preserving sensitive anaerobic bacteria alive for later culture recovery. Here we introduce a novel microbiome preservation kit (BIOME-Preserve) that facilitates recovery of anaerobic organisms from human stool held at room temperature. Using a combination of culture recovery and shallow whole-genome shotgun sequencing, we characterized the culturable anaerobes from fresh human stool and from human stool held in BIOME-Preserve for up to 120 hours. We recovered several species of interest to microbiome researchers, including *Bifidobacterium spp., Bacteroides spp., Blautia spp., Eubacterium halii, Akkermansia muciniphila*, and *Faecalibacterium prausnitzii*. Together, our results suggest BIOME-Preserve is practical for the collection, transport, and culture of anaerobic bacteria from human samples and can help provide the foundation for culture collections that can be used in further research and in the development of microbiome-based therapeutics.

**Importance:** Sequencing-based protocols for studying the human microbiome have unearthed a wealth of information about the relationship between the microbiome and human health. But these microbes cannot be leveraged as therapeutic targets without culture-based studies to phenotype species of interest and to establish culture collections for use in animal models. Contrary to popular opinion, most gastrointestinal bacteria can be cultured, yet most sample collection strategies are optimized for the preservation of nucleic acids and/or metabolites only and do not take into account considerations for preserving oxygen-sensitive anaerobes and facultative anaerobes, which comprise the majority of the human gut microbiome. A human microbiome sample transport and preservation medium such as the one described here can play an important role in enabling researchers to better understand the link between the microbiome and human health and how to leverage that link through novel microbiome-based therapeutics.

## Introduction

The human microbiome plays a fundamental role in health and disease. Most bacteria are found in the gastrointestinal (GI) tract and are primarily anaerobic and facultatively anaerobic microorganisms. A large portion of the research and analysis of the microbiome to date is based around 16S rRNA gene sequencing (1), and more recently whole genome shotgun (WGS) metagenomics sequencing, of stool samples as a proxy for the human GI microbiome (2). Such protocols were originally developed due to limitations of culture-based techniques for capturing the full microbial diversity in a sample, with some claiming that up to 99% of known microbial diversity is unculturable (3, 4). While this is true of certain environments, recent studies have shown that much of the bacterial diversity present in human stool can indeed be cultured. Lau *et al*.(5) demonstrated that the majority of bacterial taxa detected via 16S rRNA gene sequencing can be cultured via a combination of culture conditions while Ito *et al*.(6) reported successful growth of eight “unculturable” species using commercially available culture media. Similarly, Brown *et al*. isolated in pure culture 137 species from characterized and novel families, genera, and species.(7)

An increasing number of research groups are turning to culture-based approaches to complement sequencing-based protocols. Organisms isolated and phenotyped in the laboratory can be used to develop and test animal models of disease or may comprise organism collections that can serve as the basis for further research or in the development of microbial-based treatments. To fully understand and leverage the potential beneficial effects of the human microbiome as a disease treatment or health-boosting strategy, it is necessary to characterize living organisms.

The collection and successful culture of organisms from samples such as human stool is challenging and is not fully supported by current sample collection and preservation techniques. Obligate anaerobic organisms contained in the samples are rapidly killed in the presence of oxygen. Furthermore, accumulation of toxic metabolic byproducts including short chain acids and alcohols can kill microorganisms in the sample if metabolism and growth continues after collection. The growth and proliferation of some fast-growing organisms in a sample can outcompete and prevent the recovery or isolation of other more fastidious and slow growing organisms.

A fresh sample is considered the gold standard, as collecting samples in a laboratory or clinical setting followed by immediate processing can increase recovery of live organisms. However, this approach is often impractical, as it limits the population for sample collection to those in the immediate vicinity of the laboratory or clinic and with the ability to provide a sample at the time it is needed. Samples can also be frozen upon collection, which stops the metabolic process and preserves the organisms in stasis (8), but freezing organisms without a form of cryoprotectant may damage cell membranes, and samples are typically collected from the general population outside of the laboratory where appropriate freezing may not be feasible or convenient. Furthermore, shipment of frozen samples is more expensive and can be prohibitive.

These challenges can be alleviated through a sample collection device designed specifically to preserve anaerobic organisms present in a sample for culture recovery. This collection system should limit exposure to oxygen and its byproducts, prevent organism overgrowth, have practical storage and shelf life requirements, and preserve organisms for recovery even when stored at ambient temperatures.

While there have been several advances in preservation methods for DNA, RNA, and metabolites (9), which often involve agents that kill microorganisms, there are few options for preserving anaerobic bacteria in the microbiome for culture recovery. We have developed a microbiome preservation kit to address this unmet need. Using WGS sequencing and analysis of stool samples and cultured bacterial communities, we demonstrate the efficacy of this novel microbiome preservation kit, BIOME-Preserve, to preserve viability of human stool bacteria (including low abundance species) for up to five days. We suggest that BIOME-Preserve is a viable solution for microbiome research laboratories in both academia and industry to preserve viability of anaerobic bacterial species during transport and for culture-based phenotypic studies.

## Methods and Materials

### Human subjects

Three adult (18+ years of age) human donors volunteered and consented to provide a single whole stool sample for this study. The donors were asked to provide a sample using a commode specimen collection bucket (02-544-208, Fisher Scientific, Hampton, NH) and to bring the sample to the Anaerobe Systems laboratory as soon as possible after collection. The samples were immediately processed upon arrival. Donors included one adult female in her 30s, one adult male in his 70s, and one adult male in his 40s with a verbally self-reported prior recurrent *C. difficile* infection. All donors self-reported verbally no active gastrointestinal infections or recent antibiotic usage at the time of sample collection. No identifying information were stored except for the donors’ names and signatures on the consent forms. All experimental procedures were approved by Aspire IRB (Santee, CA; protocol # Anaerobe01).

### Stool preprocessing

The donor stool sample was weighed and then moved into an anaerobic chamber (AS-580, Anaerobe Systems, Morgan Hill, CA) containing a gas mixture of 4.6% CO_2_ / 5% H_2_ / 90.4% N_2_ as soon as possible after collection (within 30 minutes for donor 1, 60 minutes for donor 3, and 4 hours for donor 2). Inside the anaerobic chamber, the stool was transferred to a Ninja 400-watt blender (QB900B Shark Ninja, Needham, MA). Chilled (4°C) and pre-reduced anaerobically sterilized (PRAS) dilution blank medium (AS-9183, Anaerobe Systems) was added at a volume of 2:1 (2mL per gram) of the stool weight. The sample was blended until fully homogenized (approximately 4 minutes). Three 0.5mL aliquots of homogenized stool were collected into 2mL cryovials (#431386, Corning Life Sciences, Corning, NY) and immediately frozen at −80°C. One cryovial was sent for WGS sequencing and analysis, the others were maintained in frozen storage.

A sequential dilution of homogenized stool into PRAS Yeast Casitone Fatty Acid with Carbohydrate (YCFAC) broth (AS-680, Anaerobe Systems) was performed to obtain a 10^-5^ dilution, which was used for high density growth and in serial dilutions for low density growth on solid media as described below.

### Assessment of BIOME-Preserve microbial toxicity

A T_0_ BIOME-Preserve control tube (see **Figure 1**) was inoculated after homogenizing fresh stool to assess inhibitory effects of the components contained in the BIOME-Preserve medium on the viability of culturable organisms from stool. Homogenized stool (1.0 mL) was added directly to a single BIOME-Preserve tube (9 mL) then held at room temperature for 1 hour. Inoculation and holding were performed inside of an anaerobic chamber to eliminate any confounding effects due to oxygen.

**Figure 1.**
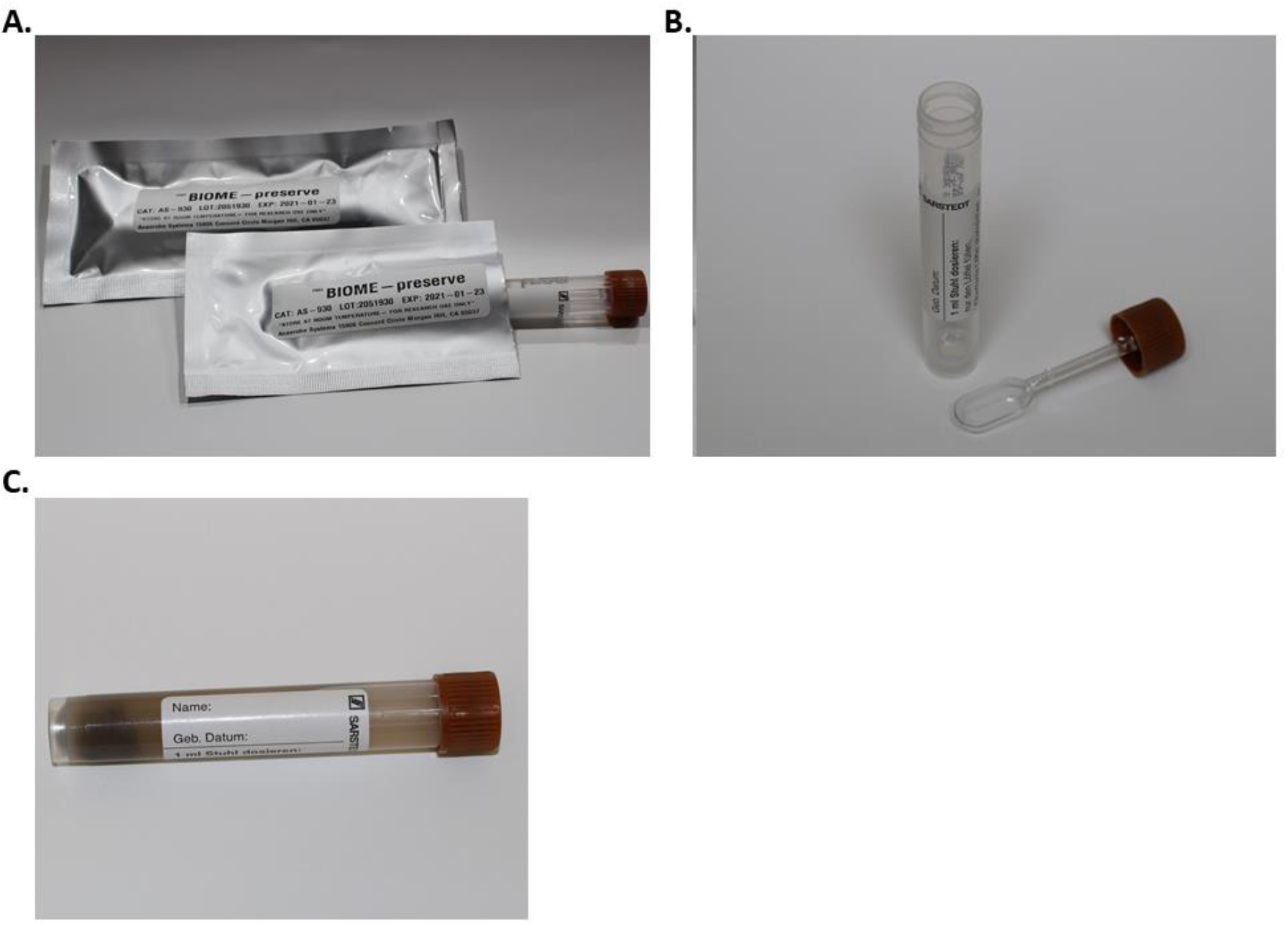
BIOME-Preserve tube.

**Figure 2.**
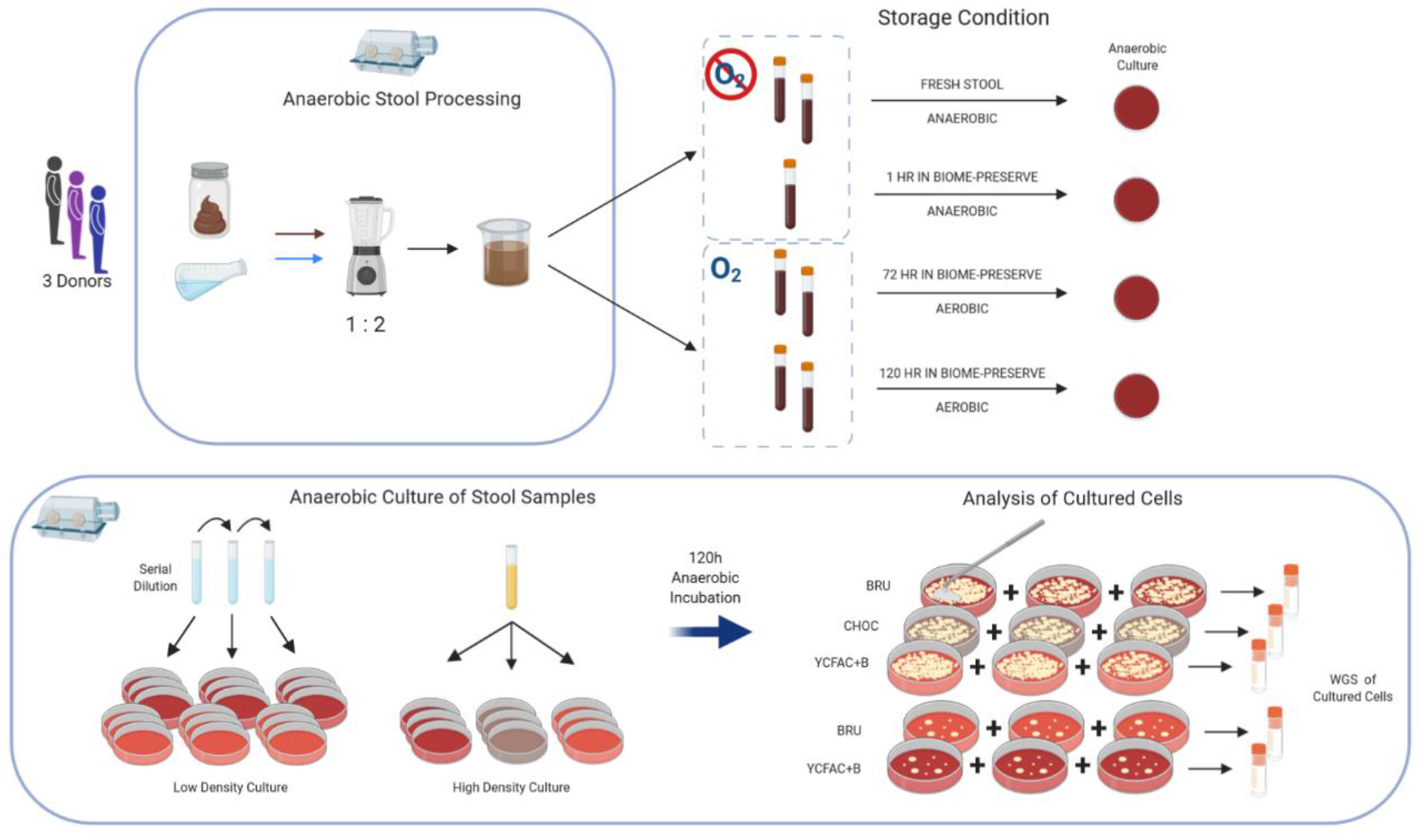
Schematic of stool collection, BIOME-Preserve inoculation, and culture of homogenized stool and BIOME-Preserve tube contents for subsequent WGS sequencing and analysis for species identification. This process was repeated 3 times for each of the 3 stool donors.

After one hour, a sequential dilution of the T_0_ BIOME-Preserve tube into YCFAC broth was performed. The 10^-5^ dilution was then used for high-density and low-density culturing on solid media as described below. A single replicate set of media were inoculated for the T_0_ BIOME-Preserve control sample as follows: 100uL of the 10^-5^ dilution were streaked onto PRAS Brucella Blood Agar (BRU, AS-141, Anaerobe Systems) and PRAS Yeast Casitone Fatty Acid with Carbohydrate and Blood (YCFAC-B, AS-677) agar plates.

### BIOME-Preserve tube inoculation

To test the ability of BIOME-Preserve to preserve anaerobic bacteria from fresh human stool, homogenized stool (1.0 mL) was added directly to two replicate BIOME-Preserve tubes (9.0 mL) and held at room temperature for 72 hours or 120 hours. These time points were chosen based on results from prior experimental data from a single donor showing minimal loss in culturable bacterial diversity between 72 and 120 hours (data not shown) and also to encompass the most likely scenarios for sample collection and/or mail transit time.

The tubes were inoculated outside of the anaerobic chamber to mimic traditional sample collection in air. At the end of each specified holding period, the tubes were inverted to mix contents, and a 0.5mL aliquot was collected from each tube and stored at −80°C. Then, 217μL were transferred from each tube to each of two replicate 7mL YCFAC broth tubes to create a 10^-3^ YCFAC BIOME-Preserve dilution. The 10^-3^ dilution was then used for high density growth and in serial dilutions for low density growth on solid media as described below.

### CFU determination for fresh stool and BIOME-Preserve tubes

To determine the CFU of fresh stool and for inoculating low-density growth plates, we performed serial dilution, targeting 10^-6^ – 10^-8^ dilutions. This process was performed twice to create two serial dilution replicate sets. The 10^-6^ – 10^-8^ dilutions were inoculated (100μL) onto PRAS BRU solid medium plates and PRAS YCFAC-B solid medium plates in triplicate. The plates were incubated in the anaerobic chamber at 37°C for 120 hours. Photographs were taken of each plate, and colonies were counted for the plates in the countable range of 30-300 colonies (see **Figure S1** for examples of CFU and high-density culture plates). CFUs from BIOME-Preserve tubes were assessed following the same protocol, except 10^-4^ – 10^-6^ dilutions were plated.

Bacterial colonies were carefully removed from plates with approximately 2,000 isolated colonies using the beveled edge of a cell lifter (08-100-240, Fisher Scientific, Hampton, NH) and transferred to a 2mL cryovial. Harvested cells from the three replicate plates from a single dilution replicate were combined into a single cryovial. 2mL of dilution blank were added to the cryovials, the cellular material was mixed via mechanical mixing, pipet mixing, and repeated inversion of the tube, and 1mL of material was transferred to a new 2mL cryovial. All cryovials were frozen immediately at −80°C; one of each duplicate tube was sent for WGS sequencing and analysis.

### High density culture of microbes from fresh stool and BIOME-Preserve tubes

To determine which microbes could be recovered from fresh stool using varying media types, 100μL of the 10^-5^ YCFAC stool dilution were streaked onto triplicate plates of the following solid media: YCFAC-B, BRU, and PRAS Chocolate Agar (CHOC, AS-244, Anaerobe Systems). Two replicate sets of media were inoculated. The plates were incubated anaerobically for 120 hours at 37°C. Following the procedure for the CFU plates described above, bacterial mat material was removed from the plates, transferred to 2mL cryovials, and split into two cryovials for a final suspended volume of 1mL. All cryovials were frozen immediately at −80°C; one of each duplicate cryovial was sent for WGS sequencing and analysis. Contents of BIOME-Preserve tubes were streaked for high-density culture following the same protocol, except the 10^-3^ YCFAC BIOME-Preserve dilution was used to streak plates. This dilution was chosen as compared to a 10^-5^ dilution from fresh stool to account for an expected 2-factor reduction in CFU during storage in BIOME-Preserve. We confirmed this through the CFU counts from solid medium plates inoculated with fresh stool and BIOME-Preserve tube contents.

### Shotgun sequencing and data analysis

To identify successfully cultured species, all cryovials collected as outlined above were sent to CoreBiome (St. Paul, Minnesota, USA) for DNA extraction and shallow WGS sequencing and analysis (BoosterShot). BoosterShot yields an average of 2 million reads per sample and enables species- and strain-level taxonomic classifications. Additionally, six samples from donor 1 were also selected for deep metagenomic sequencing (DepSeq, average of 20 million reads per sample).

Host DNA was removed from the sequencing data prior to delivery to Anaerobe Systems. Sequences were aligned against the CoreBiome Venti database, assigned operational taxonomic units (OTUs), and each OTU was assigned a taxonomy. OTUs were filtered according to the following protocol: the number of counts for each OTU was normalized to the OTU’s genome length; OTUs accounting for less than one-millionth of all species-level markers across the entire data set were removed. Additionally, OTUs with <0.01% of their unique genome regions and <1% of their total genome covered were removed. Finally, any samples with <10,000 reads were not included in the analysis. A rarefied OTU table (10,000 reads/sample) was used for downstream analyses. Full procedural details for BoosterShot are outlined in Hillman *et al*. 2018.(10) The same filtering process was applied to both BoosterShot and DeepSeq sequencing data.

To determine culturable species, we considered only OTUs with a taxonomic classification at the species (i.e. unknown species with genus, family, or higher classifications were not considered). Through a combination of estimated inoculum size on culture plates and an assessment of genome coverage acquired through the subset DeepSeq data, we considered any species culturable if its average relative abundance, according to BoosterShot data, across all replicate media plates was at least 0.01%.

To determine the change in relative abundance of culturable species between stool and BIOME-Preserve tubes and across time in BIOME-Preserve, we first determined which species were successfully cultured from fresh stool, and then compared each species’ relative abundance in fresh stool culture to its relative abundance in BIOME-Preserve culture to determine the percentage drop in relative abundance. Results were stratified according to fold-change in relative abundance (less than 1-fold, 1-fold, or 2-fold +).

Sequences and metadata are available from the European Nucleotide Database under accession number PRJEB40933.

## Results

In this study, we cultured anaerobic bacteria from fresh stool as well as stool inoculated and held in BIOME-Preserve for 72 hours and for 120 hours from three different human donors. We used three different culture media to capture a range of bacterial diversity, from highly abundant species to rarer, slow growing and/or fastidious microorganisms. Using serial dilutions, we plated high density and low-density growth culture plates, with low density culture plates used to target slow growing species that are easily outcompeted. Shallow whole genome shotgun sequencing was used to identify species that were successfully cultured, and which were impacted by storage in BIOME-Preserve.

### Culturable bacteria from fresh stool

Because of the inherent variability between individual people, we collected samples from three different adult donors, male and female, from 30-70 years of age. Although all donors were healthy (self-reported) at the time of sample collection, one donor had experienced a recurrent *Clostridioides difficile* infection a few years prior to the study, and reportedly continued to experience gastrointestinal issues believed by the donor to be related to the previous infection. No medical records were accessed or used to confirm any donor’s health status.

As described in the methods section above, we utilized DeepSeq data to validate our chosen relative abundance cut-off from BoosterShot data for identifying anaerobes recovered on culture media. We assessed the percent whole genome and unique genomic regions coverage, per DeepSeq, of all species with a relative abundance of 0.01%, per BoosterShot, in culture material collected from media plates. As Tables S1-S3 illustrate, the genomes of at least one strain per each of these species were sufficiently covered to consider those species culturable (we used CoreBiome’s coverage cutoffs for including an OTU in the OTU table as a guide: 1% for whole genome coverage and 0.01% for unique region coverage, though most strains were well above those cutoffs).

Using three different culture media and two dilution factors, we successfully cultured, from fresh stool, 173 species from donor 1, 168 species from donor 2, and 168 species from donor 3. Importantly, several of these species are of interest to microbiome researchers, including several *Bifidobacterium* and *Bacteroides* species, *Blautia* species, *Akkermansia muciniphila*, and *Faecalibacterium prausnitzii*.

Comparing sequencing data only, we observed three unique bacterial communities in terms of diversity and composition **(Figure 3)**. All three donors were dominated by species in the Firmicutes phylum (53.5% for donor 1, 62.7% for donor 2, and 67.2% for donor 3, see **Figure S2**), while all had a similar relative abundances of Bacteroidetes. Proteobacteria were detected at 2.3% and 1.3% relative abundance in donor 2 and donor 3, respectively, but less than 1% relative abundance in donor 1. Notably, Verrucomicrobia were detected only in donor 1. The diversity of species present (see tables S4-S6) across the donors provided a good foundation to test the ability of BIOME-Preserve to preserve a variety of bacterial species across different individual donors.

**Figure 3:**
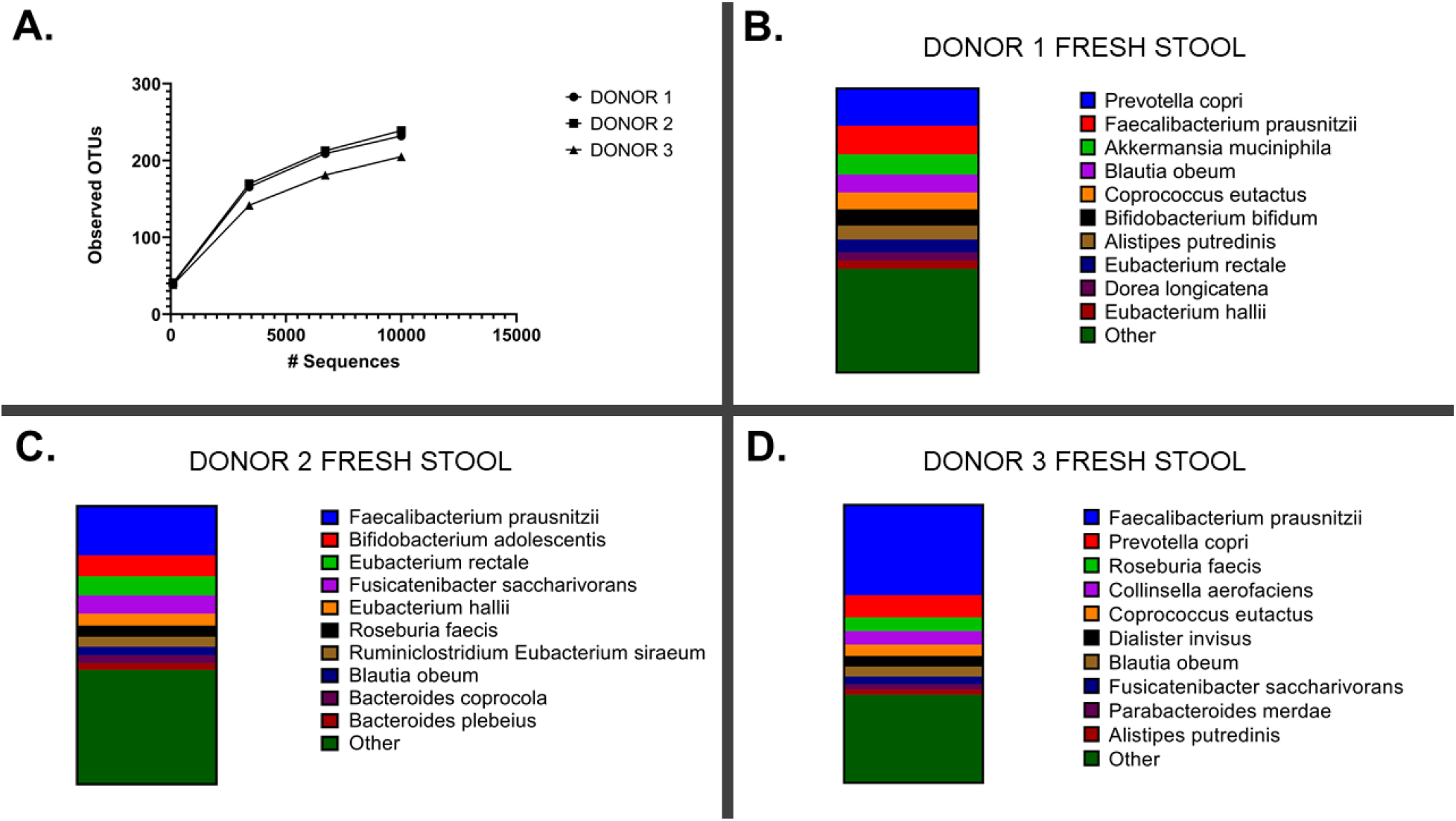
Shallow WGS sequencing of homogenized stool contents reveals a variety of bacterial species present across the three human donors in this study. **A)** Rarefaction curve illustrating total number of observed OTUs per donor; samples rarefied to 10,000 sequences per sample max. The top ten species detected in donor 1 (**B**), donor 2 (**C**), and donor 3 (**D**), are plotted, with remaining species grouped into “other.”

### Assessment of the effect of BIOME-Preserve contents on stool microorganisms

Before assessing the capacity of BIOME-Preserve to maintain anaerobic organisms alive for culture, we first determined whether the additives in BIOME-Preserve harmed anaerobic organisms from human stool. We sequenced cultured cells from homogenized stool collected from donor 1 and from stool held in BIOME-Preserve for 1 hour. It is important to note that this tube, unlike the other BIOME-Preserve tubes, was inoculated and held under anaerobic conditions to eliminate any harmful effects of temporary exposure to oxygen and ensure the only variables tested were the BIOME-Preserve additives.

As seen in **Figure 4**, there was minimal difference in the total number of cultured species from the BIOME-Preserve tubes held for 1 hour compared to fresh homogenized stool. A total of 163 species were recovered on culture media from the T_0_ BIOME-Preserve tube, compared to 173 species recovered from fresh, homogenized stool. A similar proportion of high, middle, and low-abundance species in fresh stool (as determined via WGS) were recovered from fresh stool culture and T_0_ BIOME-Preserve culture.

**Figure 4.**
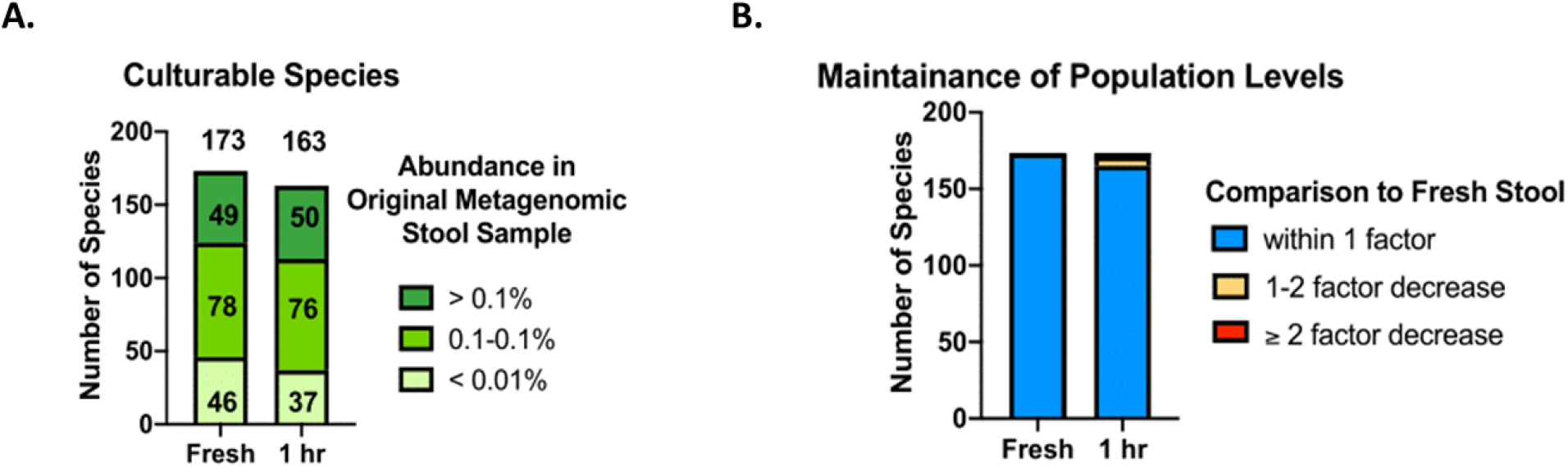
**A)** The number of species recovered from fresh stool and the T_0_ BIOME-Preserve tube, stratified by relative abundance in fresh stool. **B)** The relative abundance of species cultured both from fresh stool and from the BIOME-Preserve T_0_ tube, stratified by change in relative abundance. The majority of culturable species had less than a 1-factor drop in relative abundance.

### Effects of holding time in BIOME-Preserve on culturable anaerobes

We next determined whether inoculation outside of the anaerobic chamber (i.e., in the regular laboratory environment) and longer holding times had any effect on the anaerobes that could be cultured from BIOME-Preserve compared to anaerobes that could be cultured from fresh stool. We held BIOME-Preserve tubes inoculated with homogenized stool at room temperature for two time periods (72 hours and 120 hours), chosen for relevance to typical lengths of time it takes for samples to arrive in the laboratory from the field, or to transit from one location to the next for processing.

Sequencing of culture material from solid media revealed that we cultured 163 and 183 species from donor 1 samples held in BIOME-Preserve for 72 and 120 hours, respectively, 145 and 133 from donor 2, and 144 and 139 from donor 3 **(Figure 5A)**. We recovered from BIOME-Preserve species at both high and very low abundances in fresh stool (**Figure 5B**), as well as a small number of species that we did not recover from fresh stool.

**Figure 5.**
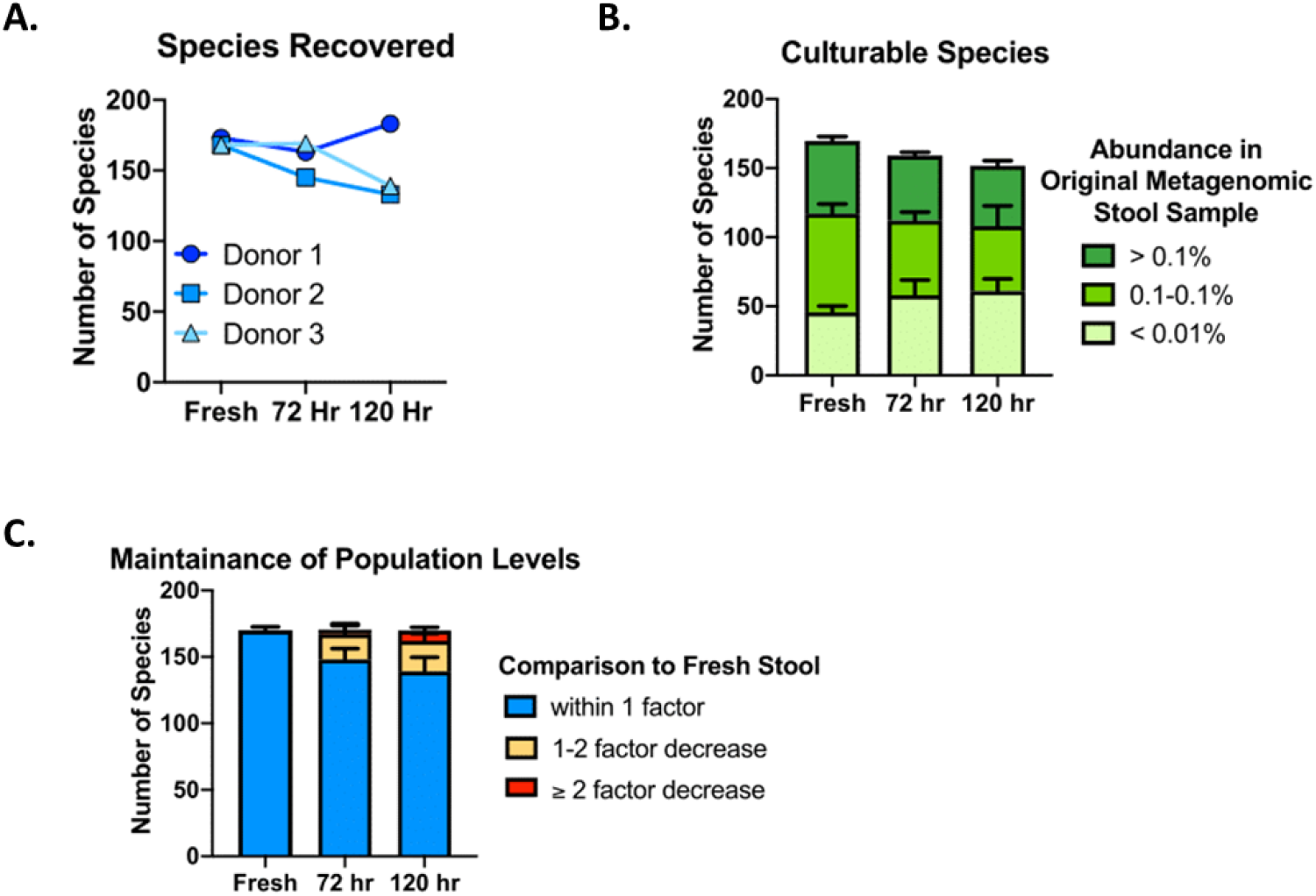
**A)** The number of species recovered from fresh human stool and from BIOME-Preserve after 72 and 120 hours. **B)** Species cultured from BIOME-Preserve tubes stratified by relative abundance in fresh stool. **C)** Species culturable from stool that were also cultured from BIOME-Preserve tubes after 72 hours and 120 hours, stratified by fold-change in relative abundance. The majority of culturable species had less than a 1-factor drop in relative abundance.

Considering only those species that were cultured both from fresh stool and from BIOME-Preserve samples, we cultured 88% of culturable species in stool from BIOME-Preserve after 72 hours and 87% after 120 hours for donor 1. Those numbers were 83% and 77% for donor 2, and 89% and 81% for donor 3 **(Figure 5C)**. Across all donors, we cultured an average of 87% of culturable species from stool after 72 hours in BIOME-Preserve, and 82% after 120 hours.

As expected, we observed medium-dependent recovery of bacterial species, with CHOC and BRU agar yielding more similar microbial communities (**Figure 6**). YCFAC-B agar performed the poorest in terms of the number of species it was able to recover, but it was able to recover specific organisms that CHOC and BRU agar could not (**Tables S7-8**). These results support the need for multiple media types to recover the majority of bacterial diversity present in human stool.

**Figure 6.**
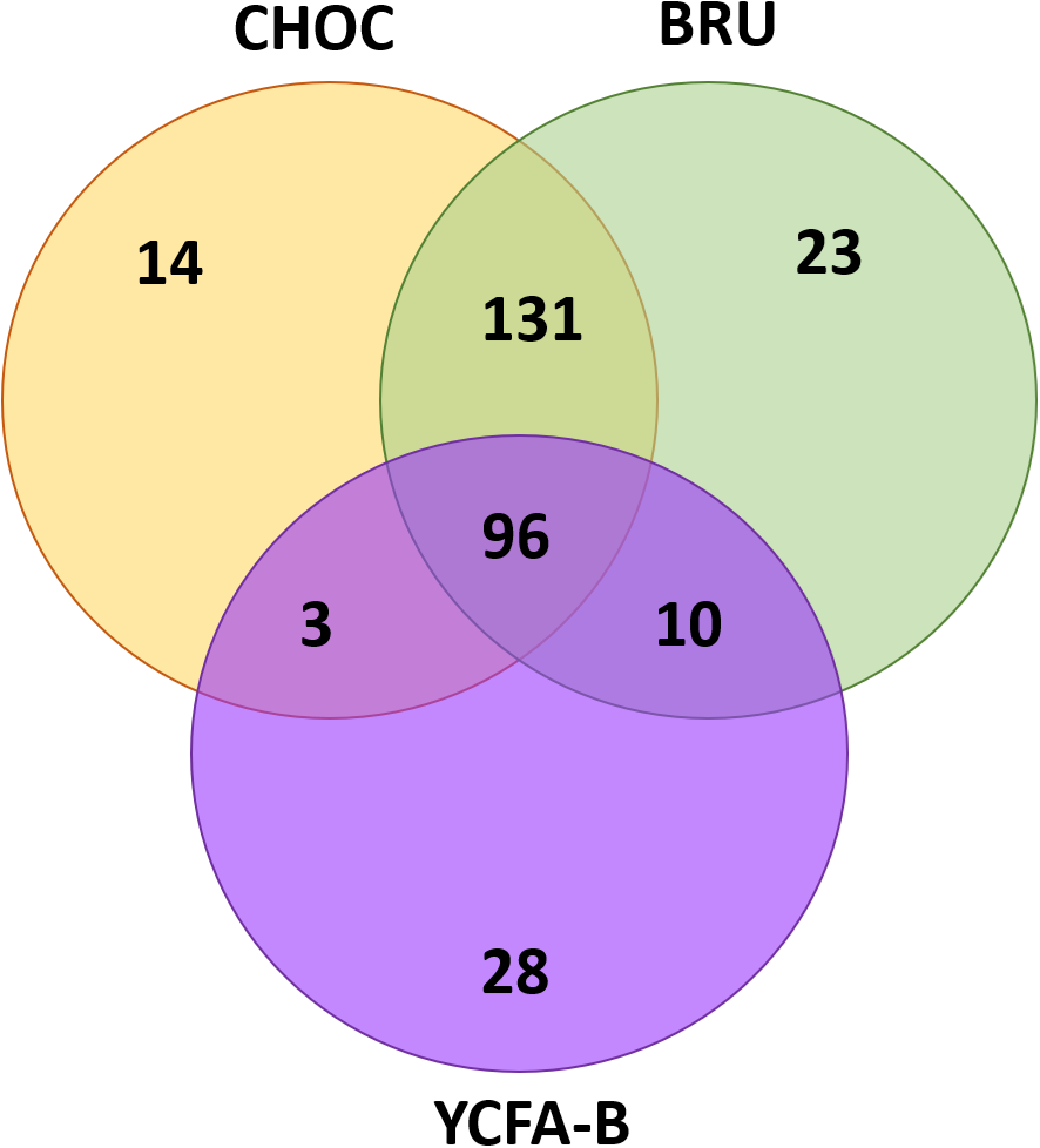
Venn diagram illustrating media-specific growth of anaerobic bacteria. CHOC and BRU solid media captured many of the same species; YCFAC-B solid medium facilitated growth of a different set of bacterial species.

Because we plated low-density plates in addition to high density plates to increase our chances of culturing easily outcompeted slow-growing organisms and lower abundance species, we next assessed which organisms were more easily cultured on low-density plates. Several species had a greater than 100% increase in relative abundance on low-density plates compared to the high-density plates of the same media type. Most of these organisms fit into seven genera: *Bacteroides, Collinsella, Lachnoclostridium clostridium, Roseburia*, and *Streptococcus*, although several species belonging to other genera were also captured, including *Akkermansia muciniphila* (see **Table 1**).

**Table 1.**
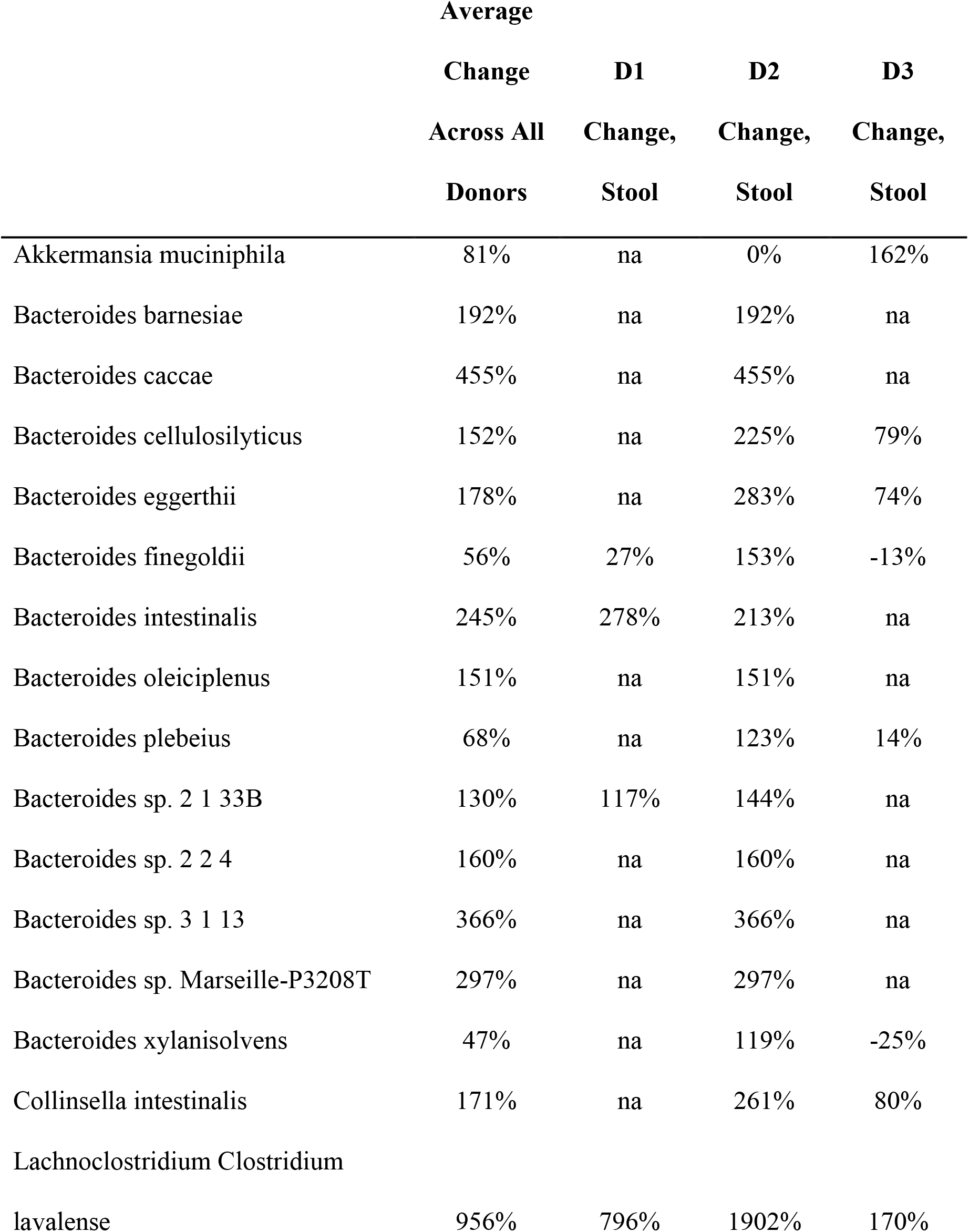
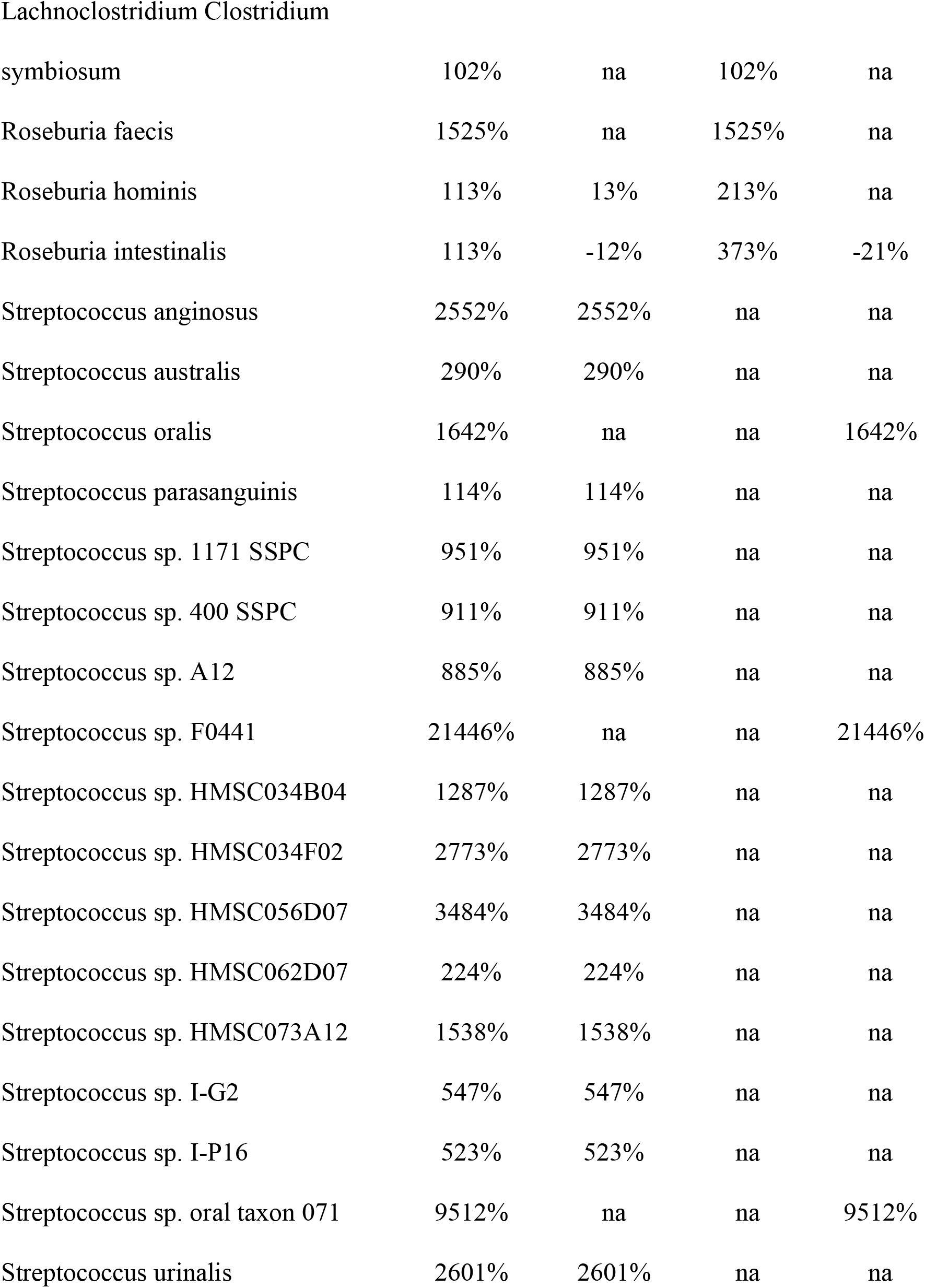
Percent change in relative abundances of species more successfully recovered on low density plates compared to high density plates (species recovered from fresh stool).

### Effects of ultra-cold storage on BIOME-Preserve

Because many microbiome studies do not process samples immediately for nucleic acids extraction, metabolomics, or culture setup, we compared the growth of anaerobes cultured from BIOME-Preserve held at room temperature to the growth of anaerobes cultured from BIOME-Preserve held at room temperature and then frozen at −80°C for one week. Using homogenized stool previously provided by donor 1, we demonstrated that over 90% of the anaerobes cultured from unfrozen BIOME-Preserve were also cultured from BIOME-Preserve stored at −80°C (**Figure 7A**). Importantly, a single freeze-thaw cycle did not hinder our ability to culture organisms at very low abundance in fresh stool from the BIOME-Preserve samples (**Figure 7B**).

## Discussion

Here we demonstrate that BIOME-Preserve is an effective method for preserving anaerobic bacteria for culture recovery for up to 5 days. From stool samples of 3 donors and using three media types and two dilution factors, we cultured from fresh stool an average of 170 species detected via shallow WGS sequencing. After 72 hours in BIOME-Preserve, we cultured an average of 150 species, and after 120 hours in BIOME-Preserve we also cultured an average of 150 species; at both time points we cultured a small number of species that were not recovered from fresh stool.

Of the species cultured from fresh stool and BIOME-Preserve samples, 87% were minimally impacted by 72 hours in BIOME-Preserve; this number dropped to 82% after 120 hours. Because the BIOME-Preserve tubes were inoculated, capped, and held outside of an anaerobic chamber, these numbers are reflective of expected results given typical laboratory procedures. Further protection may be afforded if stool is added to BIOME-Preserve inside of an anaerobic chamber. Because such an approach is not always feasible, especially during field studies, we only assessed the efficacy of BIOME-Preserve in the most likely real-world scenario.

The change in relative abundance between fresh stool and BIOME-Preserve was within 1-fold for the vast majority of anaerobic species that we cultured both from stool and BIOME-Preserve, with a very small number experiencing a 2-fold change in diversity. Considering the average human gut contains 10^12^ microorganisms, a 2-fold change in relative abundance still equates to large numbers of bacterial cells and demonstrates that BIOME-Preserve is much more stringent for maintaining live organisms than any other available method. For example, we cultured *Faecalibacterium prausnitzii* from both stool and BIOME-Preserve with a 1-fold change in relative abundance in BIOME-Preserve; by comparison, Papanicolas *et al*. (11) reported a 12fold loss in this species when stool samples were processed aerobically vs. anaerobically. This suggests that BIOME-Preserve could be an appropriate tool for preserving microbiome communities intended for therapeutic purposes such as FMT or for other downstream applications such as studies using humanized animal models. It can also substantially reduce the laboratory resources needed at collection sites.

We collected stool from three adult humans, two male and one female, one of which was elderly and one of which had a self-reported previous recurrent *C. difficile* infection, to assess the ability of BIOME-Preserve to protect a wide variety of organisms from different human individuals. While the microbial communities were notably different from one another, there were no donorspecific effects on our ability to recover organisms from fresh stool or from BIOME-Preserve. We cultured 173 species from donor 1, 168 species from donor 2, and 168 species from donor 3 with 85%, 85%, and 80% of those species experiencing a less than 1-fold difference in relative abundance after 5 days in BIOME-Preserve. With 381, 516, and 426 species detected via WGS sequencing in donor 1, 2, and 3, respectively, our protocol successfully cultured, on average, 39% of fecal species detected via sequencing. Notably, a single freeze-thaw cycle had minimal effect on the anaerobic bacteria we successfully cultured from BIOME-Preserve.

Our results, based strictly on the number of species recovered, are similar to those reported by Browne, *et al*., (7) who successfully cultured 137 species, including 45 candidate novel species using only a single medium type (YCFA). Based on percentages of what is detected in stool, we recovered 20% less than that reported by Ito *et al*. (61%), and Lau *et al*.(5) reported culture success rates over 90%. Those two groups used 27 and 33 different media, respectively, and Lau *et al*. used anaerobic and aerobic culture conditions (5); the vast number of plates utilized by these two protocols provides many more chances to obtain very low abundance species and easily outcompeted species compared to our protocol. Cultures consisted of a vast and highly diverse population with many factors such as sampling variation, plate spreading, and intercommunity growth all affecting whether or not a species could be recovered on culture media. We also did not count every species with a relative abundance >0% on culture media as successfully cultured; therefore, based on our cutoff (at least 0.01% relative abundance), some species that may have indeed grown on culture are not included in our percentages.

Additionally, it is difficult to perform direct comparisons; for example, Browne *et al*. recovered less species than we in terms of sheer number but outperformed in terms of percentage compared to stool (72%). Such variability could be due to cross-donor differences as the number of species present in a “typical” human gut can range from 500-1,000 (12, 13). Additionally, the number of species detected via sequencing may not be representative of the truly culturable proportion, as some DNA could belong to transient organisms no longer present or to dead organisms and therefore may just be artifacts. Sequencing method and taxonomy databases can also affect results; we used WGS sequencing and analysis, which provides more accurate species calls than 16S rRNA gene sequencing; however, we also eliminated any OTUs that did not receive a confident species-level identification. While these OTUs do represent individual species, we found it more valuable to the community to provide information on known species at this time; had we included these OTUs in our final counts our final numbers may have been more comparable to those achieved by other groups.

Based on results from other groups, we chose three media types to maximize the culturable bacterial diversity: Brucella blood agar, Chocolate agar, and Yeast Casitone Fatty Acid agar supplemented with carbohydrates and blood. The most abundant species present in all three donors were easily cultured using these three media types. We also successfully cultured several anaerobes that were at very low abundance in the original stool samples, as well as several species of interest to the microbiome research community. Notably, two species of interest (*Prevotella copri* and *Faecalibacterium prausnitzii*) were recovered at a higher relative abundance on low density plates than on high density plates, suggesting that more dilute samples may be necessary to capture slow growers and easily outcompeted organisms.

Brucella blood agar and Chocolate agar yielded similar microbial communities on culture. Both successfully recovered several members of the *Bacteroides* genus that were not well-recovered using YCFAC-B agar. This includes *Bacteroides fragilis*, which has recently been dubbed a “next-generation” probiotic (14) and has been tested in the context of several disease states, including autism spectrum disorder (15). On the other hand, YCFAC-B was particularly useful compared to BRU and CHOC agar for recovering a wide range of common human anaerobes, including the butyrate producers *Eubacterium rectale* and *Faecalibacterium prausnitzii*, a major component of the healthy human microbiome and an important contributor to intestinal health (16–18). YCFAC-B also enabled recovery of several low-abundance (<0.01%) *Streptococcus* and *Lactobacillus* species, both from fresh stool and from BIOME-Preserve samples. All three media types recovered several *Bifidobacterium* and *Blautia* species; some *Blautia* recovered better on YCFAC-B while others did better on BRU or CHOC. *Blautia* is of increasing interest to the research community due to its inverse association with obesity, visceral fat, disturbances in glucose metabolism, and intestinal inflammation (19, 20).

Of note, while we successfully cultured several *Prevotella* species from fresh stool on all three media types, we struggled to culture one species of great interest to the microbiome research community, *Prevotella copri*, from BIOME-Preserve samples, only capturing it at one time point (72 hour BIOME-Preserve) from a single donor (donor 3) and at very low relative abundance (0.07%). The current experimental methods were not sufficient to determine confidently whether these results are due to toxicity from BIOME-Preserve or metabolic byproducts produced by other anaerobes in the community; however, the fact that we could recover the organism from fresh stool and from the T_0_ BIOME-Preserve tube (albeit at a much lower abundance than from stool) suggests that something native to BIOME-Preserve or to community dynamics occurring over the span of days impacted this organism. It is possible that different media choice, different dilution factor(s), and/or different culture protocols may have successfully recovered this organism.

Recently, Martinez *et al*. (21) reported a sample collection and transport device (GutAlive) designed to maintain viability of extremely oxygen sensitive (EOS) bacteria common in human stool. They report higher bacterial diversity from GutAlive than traditional stool collection devices, with losses in diversity observed after 24-48 hours, demonstrating that GutAlive is a better approach to conventional stool collection devices. Because Martinez *et al*. did not compare either stool collection device to the bacterial community of fresh stool, their protocol was unable to determine the extent to which GutAlive can preserve whole stool communities and there is no way to determine the true magnitude of loss of anaerobic species using GutAlive. In contrast, we demonstrated that nearly 85% of culturable species from fresh stool remain culturable after 5 days in BIOME-Preserve, suggesting that not only does a collection device specifically tailored for anaerobes perform better than traditional stool collection devices as Martinez *et al*. showed, but also demonstrating that BIOME-Preserve specifically is an attractive option for collecting, storing, and recovering the majority of anaerobes that can also be cultured from human stool.

Unlike other similar studies, we did not protect the original fecal sample from oxygen between time of production until it arrived in the anaerobic chamber for homogenization, which may have some detrimental effects on extremely oxygen sensitive organisms. Notably, while the samples from donors 1 and 3 were transferred to the anaerobic chamber within 1 hour, the sample from donor 2 was not transferred to the anaerobic chamber until 4 hours after it was produced. Assessing extremely oxygen sensitive species (EOS) (22–24), we found that we successfully cultured several, including *Clostridium phoeensis* (which was actually recovered at a higher relative abundance from cultured BIOME-Preserve samples than from cultured stool) and *F. prausnitzii* (18, 25). Other EOS were not successfully recovered, but as they were already low abundance in fresh stool (<0.05%) it is difficult to determine whether failure to culture is a result of oxygen damage or increased difficulty in capturing very low abundance organisms from a diverse microbial community. Our data suggest that our experimental procedures prior to anaerobic chamber work did not pose a significant detriment to our ability to capture anaerobic organisms generally; nevertheless, we recommend that for the best results, fecal material be transferred to BIOME-Preserve tubes as soon as possible after collection or otherwise protected from oxygen if immediate transfer to BIOME-Preserve isn’t possible.

Altogether, our results show that BIOME-Preserve is an effective tool for collecting and preserving human stool samples for culture recovery of anaerobic organisms in the laboratory. The ability of BIOME-Preserve to preserve over 85% of culturable diversity after five days at room temperature demonstrates the utility of this project for any microbiome study unable to immediately freeze or process samples, and will likely facilitate studies that until now have been infeasible.

## Acknowledgements

We thank the scientific team at CoreBiome for experimental design and data analysis support. We also thank our study volunteers.

This study was funded by Anaerobe Systems, Morgan Hill, CA. H.L. is an employee of Anaerobe Systems. S.M.C. is president of Anaerobe Systems. E.H. is an independent research consultant.

